# Connections across regional glymphatic clearance, neural activity and amyloid-β deposition in cortex

**DOI:** 10.64898/2026.04.23.720377

**Authors:** Yifei Li, Xiao Zhu, Ying Zhou, Xuting Zhang, Ziyu Zhou, Kai Wei, Jianzhong Sun, Min Lou

## Abstract

Neural activity inevitably produces waste, which promotes neurodegeneration with topographic features. The glymphatic system is important for waste clearance. However, the spatial characteristics of glymphatic clearance across cortex, the pattern it coupled with neural activity, and whether those mismatch regions overlap with known amyloid-prone cortical regions in human remain unexplored. Here, by intrathecal administration of gadolinium-based contrast agents, glymphatic influx and clearance patterns across cortex in 96 participants are depicted via Glymphatic MRI. Analyses integrating public post-mortem transcriptomic profiles from Allen Human Brain Atlas indirectly indicate that, genes related with excitatory and inhibitory neurons, and pathways engaging in synaptic function were spatially enriched in regions with faster glymphatic clearance. Fractional amplitude of low frequency fluctuation was calculated from resting-state fMRI to represent neural activity. At the regional level, based on a subgroup with rs-fMRI (N = 15), regional glymphatic clearance was positively coupled with spontaneous neural activity. The mismatch index, reflecting decoupling between spontaneous neural activity and glymphatic clearance function, showed positive spatial associations with an external group-average ^¹¹^C-PiB SUVR map. Together, this study for the first time demonstrates that neural activity is associated with glymphatic dynamics from transcriptional to physiological level. The mismatch magnitude between these two processes may potentiate regional vulnerability to proteopathy and subsequent neurodegeneration in cortex.

## Introduction

The accumulation of metabolic waste serves as a leading cause in multiple neurological diseases^1^. The brain’s remarkable neurocomputational capacity, supported by synchronized neuronal activity, inherently generates substantial metabolic waste^2^. In the last few decades, evidence suggested that higher regional activity is associated with increased susceptibility to pathological hallmarks such as amyloid-β (Aβ) deposition^3, 4^. While prevailing hypotheses predominantly emphasize production side, waste deposition could equally be attributed to the clearance side. The equilibrium disruption between waste generation and elimination may ultimately manifest as regional vulnerability and waste deposition in neurodegenerative diseases.

The landmark characterization of the glymphatic system has reshaped our understanding of cerebral waste clearance homeostasis. In this system, cerebrospinal fluid (CSF) flows into the brain via perivascular space (i.e., influx), enters the interstitial compartment through an astrocytic AQP4-dependent process, and finally flushes out metabolic waste, including Aβ and tau (i.e., clearance)^5, 6^. Compelling experimental evidence demonstrates that glymphatic failure precedes and potentiates Aβ pathogenesis, with genetic ablation of AQP4 polarization exacerbating both amyloidosis and cognitive decline in AD models^7^. Crucially, the impairment of glymphatic clearance is not temporally and spatially uniform. In aging rodents, widespread astrogliosis and loss of perivascular AQP4 polarization is evident especially within the cortex^8^, but the regional glymphatic dynamics remain incompletely characterized. How and to what extent regional glymphatic dynamics contribute to Aβ accumulation in human is largely unknown.

Several evidence suggests neuronal activity as a dual-edged modulator of cerebral homeostasis. While driving metabolic waste production, it may simultaneously activate compensatory clearance. Better glymphatic clearance significantly correlated with higher FDG-PET uptake^9^, an indicator of metabolic demand and neural activity. Fultz et al first pointed that neural and hemodynamic rhythms are coupled with CSF dynamics in human^10^. Recently, three independent groups demonstrated that neuronal oscillations serve as an important organizer of brain clearance^11, 12^, possibly through neurovascular coupling^13^. Together, these findings reshaped our understanding of the relationship between glymphatic dynamics and neural activity^14^, and inspired us to systematically investigate their spatiotemporal coordination.

Topographically patterned vulnerability to waste deposition is a key feature in neurodegeneration. We hypothesized that regional vulnerability is dictated not solely by waste production due to local neural activity, but also by the magnitude of mismatch between neural activity and the clearance capacity. To test this hypothesis, we systematically followed three steps: 1) Employing Glymphatic MRI by intrathecal administration of gadolinium-based contrast agents^15^, we quantified spatiotemporally resolved glymphatic dynamics across cortex. Following injection, the tracer flows from the perivascular space into the brain parenchyma. Thus, the early phase of cortical signal enhancement (e.g., at 4.5 h post-injection) primarily reflects tracer influx. As the tracer gradually eliminated, prolonged residual signal at a delayed time point (e.g., at 39 h post-injection) indicates abnormal tracer retention, serving as a proxy for impaired clearance. To minimize individual variability and capture macroscale axis of variance in cortical glymphatic features, gradients of glymphatic influx and clearance were calculated, organizing regions along spatially continuous spectra^16, 17^. 2) Integrating with post-mortem transcriptomic profiles^18^, we tested whether transcriptomic features of glymphatic dynamics were related with components of neural activity. Then, in a subgroup with additional resting-state functional MRI (rs-fMRI), we examined whether glymphatic clearance coupled with regional neural activity and further defined neural activity-glymphatic clearance mismatch regions. 3) Leveraging open-source ^11^C-PiB PET SUVR dataset^19^, we tested whether mismatch regions overlap with amyloid-prone cortical regions. Study pipelines are shown in Fig. 1.

**Fig. 1.**
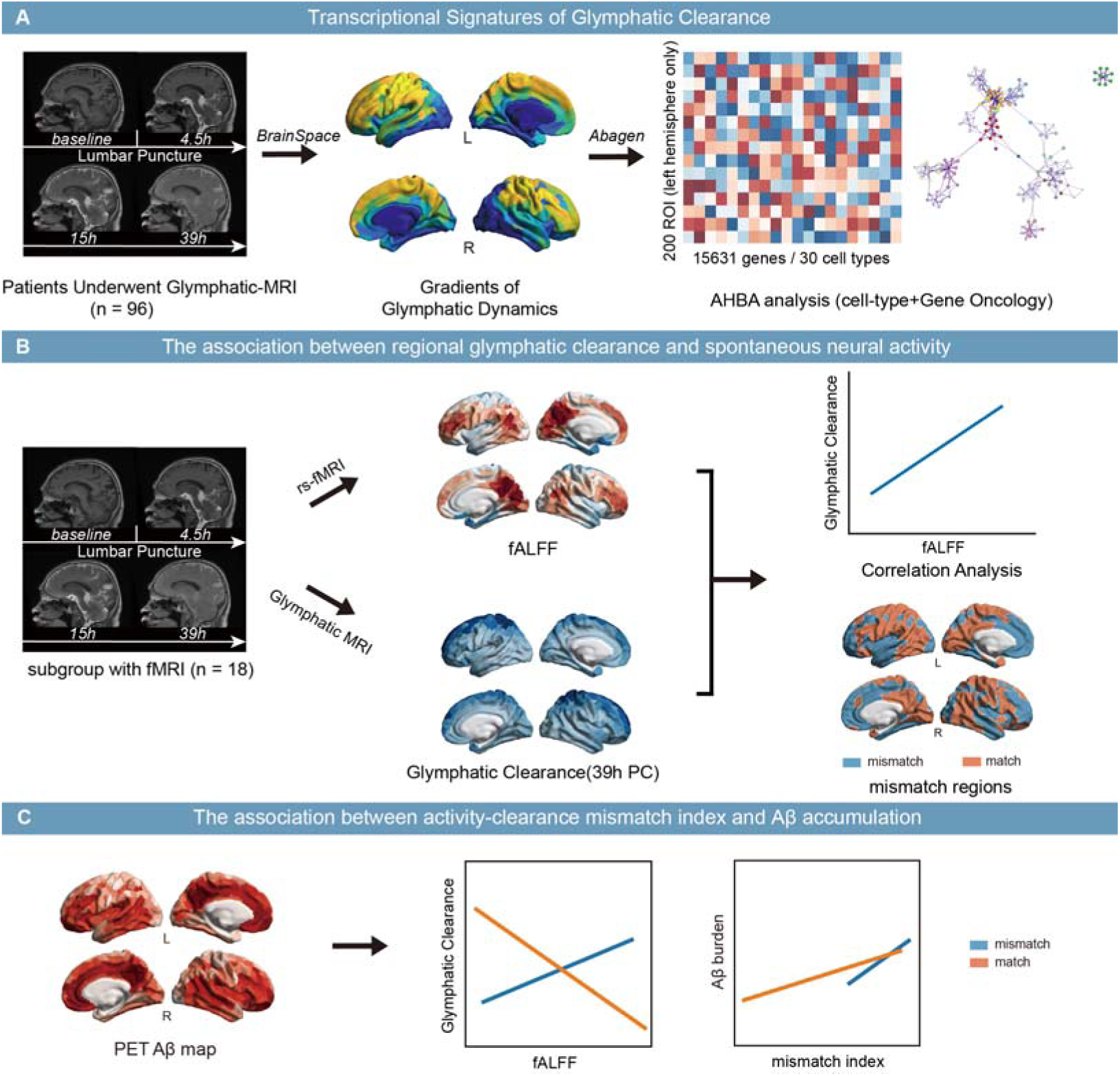
Overview for analysis pipeline. (A) The pipeline for computing glymphatic dynamics and transcriptional analysis. Gradients of cortical glymphatic influx and clearance were derived using BrainSpace. Utilizing Abagen toolbox, the glymphatic patterns were further analyzed with transcriptional data from Allen Human Brain Atlas (AHBA). Both gene ontological and cell type analyses were conducted. (B) The relationship between glymphatic clearance function and spontaneous neural activity measured as fALFF was explored in a subgroup with resting-state fMRI (n = 15). (C) The mismatch index between glymphatic clearance function and spontaneous neural activity was defined to investigate its association with amyloid-β (Aβ) using public dataset.

## Materials and Methods

### Patients

Written informed consent was obtained from all patients and the study was approved by the local ethic committees. Coherent with prior studies^20^, patients with indications for lumbar puncture and voluntary participation in this research were enrolled from July 2018 to December 2024. Exclusion criteria were history of a hypersensitivity reaction to contrast agents, history of severe allergic reactions in general, evidence of renal dysfunction, and pregnant or breastfeeding females. Additional exclusion criteria in the study were: (1) failure of T1 segmentation; (2) failure of signal normalization.

Participants with rs-fMRI were further included in fMRI-related analysis. Exclusion criteria for this subgroup included: (1) without fMRI; (2) excessive head motion (mean Power framewise displacement > 0.3); (3) with intracranial diseases proven to be associated with glymphatic dysfunction, including encephalitis, normal pressure hydrocephalus, and neurodegenerative diseases.

Considering diagnosis, neurodegenerative diseases with intracranial pathology included normal pressure hydrocephalus, Parkinson’s disease, possible cerebral amyloid angiopathy, multiple system atrophy, Alzheimer’s disease, and Lewy body dementia. All patients received an intrathecal injection of gadolinium at 3:30∼4:00 pm on day 1, then underwent 3D-T1 at 4.5 hours, 15 hours and 39 hours after intrathecal injection of gadolinium (Glymphatic MRI). Participants remained hospitalized for at least 39 h and were allowed to follow their usual nighttime sleep. Sleep duration and quality were not objectively monitored.

### Intrathecal administration of gadolinium

The site of intrathecal injection of the contrast agent was L3-4 or L4-5 lumbar intervertebral space. Intrathecal injection of 1ml of 0.5mmol/ml gadodiamide (Omniscan; GE Healthcare) was preceded by verifying the correct position of the syringe tip in subarachnoid space in terms of CSF backflow from the puncture needle. Following needle removal, the study subjects were instructed to rotate around the body’s long axis twice and then remained in the supine position until 4 hours after intrathecal injection.

### MRI protocol

All patients underwent MR by a 3.0T MR scanner using an 8-channel brain phased array coil. The imaging parameters of 3D-T1: repetition time =7.3 ms, echo time = 3.0 ms, flip angle = 8°, thickness = 1 mm, field of view = 25 × 25 cm^2^, matrix = 250 × 250. This sequence was acquired for four time points, i.e., before and 4.5, 15, and 39 hours after injection, to gain Glymphatic MRI. For rs-fMRI, it was acquired using a T2-weighted gradient echo-planar imaging sequence, time of repetition = 2000 ms, time of echo = 30 ms, flip angle = 77°, field of view = 240 × 240 mm^2^, matrix = 64 × 64, slice thickness = 4.0 mm, slice number = 38.

### Glymphatic MRI processing

#### Cortical glymphatic signal percentage change calculation

For Glymphatic MRI preprocessing, individual baseline T1 images were non-rigidly registered to ICBM template using FreeSurfer^21, 22^ and further served as the reference. Serial T1 images were co-registered to this reference using SPM12. Mean signal for each region of interest (ROI) from 400-parcel Schaefer (Schaefer-400) atlas^23^ was extracted for serial T1 images and then normalized by dividing the corresponding time point of vitreous body signal according to previous studies^24^, termed as signal ratio. Normalized T1 images were independently quality-checked by two neurologists (Y.L. and X.Z.). Signal percentage change (PC) for each ROI from baseline to 4.5, 15, or 39 hours was calculated according to equation (1).

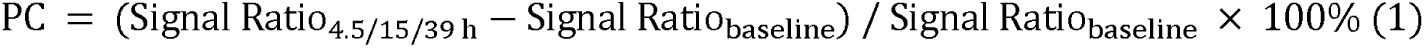

The PC at the early time point (4.5 h) was utilized as a macroscopic imaging proxy for glymphatic influx. Conversely, the PC at the delayed time point (39 h) represents prolonged tracer retention, serving as a proxy for impaired glymphatic clearance. Higher 39h PC indicates greater tracer retention, thereby reflecting worse clearance capacity.

#### Cortical glymphatic gradient calculation

Glymphatic gradients were calculated for both 4.5h and 39h. For each timepoint, PC map across cortex was used to generate a covariance matrix across participants. First, it involves z-scoring each participant’s PC map followed by z-scoring the data across all participants for each ROI based on Schaefer-400 atlas. Second, the pairwise correlations between ROIs were calculated to construct the covariance matrix. Then, BrainSpace toolbox^16^ was applied to find the gradients for each PC map. A Diffusion mapping manifold algorithm (with default parameters) was used to derive the first gradient of the covariance matrix and used for further investigation.

### Processing and analysis of functional MRI

Rs-fMRI was preprocessed by DPABI^25^. Preprocessing steps included: 1) convert the DICOM format to NIFTI format, 2) slice timing (removing the first 10 volumes), 3) realignment (to the middle volume), 4) nuisance regression (i.e., Friston-24 head motion parameters, white matter signal and cerebrospinal fluid signal which were extracted from individual T1 segmentation), 5) motion scrubbing (the volume of Power framewise displacement > 0.2 and its previous 1 and post 2 volumes were censored), 6) normalization (by DARTEL to Montreal Neurological Institute space into 3×3×3 mm voxels), 7) smoothing (with a Gaussian kernel of 6×6×6 mm), and 8) band-pass filtering (0.01-0.1 Hz). After preprocessing, fractional amplitude of low-frequency fluctuation (fALFF) and regional homogeneity (ReHo) were calculated using DPABI. Briefly, power spectrum was computed for each ROI, the square root of which is the amplitude at each frequency. The fALFF map was calculated as the sum amplitude within 0.01-0.1 Hz divided by the sum of amplitudes over the whole frequency range for each ROI. ReHo was calculated for each voxel using Kendall’s coefficient of concordance, assessing the temporal similarity with its 26 nearest voxels. The standardized ReHo was derived by normalizing each voxel’s Kendall’s coefficient of concordance against the whole brain’s mean value, followed by smoothing with a 6mm FWMH Gaussian kernel. The z-score map of fALFF was used in formal analysis, with ReHo used in sensitivity analysis.

### Transcriptomic analysis

#### Transcriptomic data

Regional microarray expression data were derived from Allen Human Brain Atlas (AHBA)^26^ based on six post-mortem brains (1 female, ages 24–57, 42.5 ± 13.38). Since the AHBA only included right hemisphere for two donors, we only considered left hemisphere for further analysis as published before^18^. Abagen toolbox was used to process and map the transcriptomic data onto the Schaefer-400 atlas based on previous studies^18, 27^. Briefly, it included pre-processing and assigning tissue samples to ROIs of atlas, which was based on guidance provided by Arnatkeviciute et al.^18^. As for pre-processing, it included gene information re-annotation, data filtering, and probe selection. This procedure retained 15631 probes, each representing a unique gene. As for assignment, sample-to-region matching was constrained by hemisphere and cortical-subcortical divisions. Due to the number and site mismatch of probes and ROIs, there would be regions failed to be assigned with a value. To ensure each ROI was assigned a value, the sample closest to the region was selected to represent it (i.e., “Centroid”). This procedure resulted in a 200 × 15631 matrix, denoting the expression of 15631 genes across 200 nodes.

#### Association between neuroimaging and transcriptomic data

We calculated partial least squares (PLS) regression between cortical glymphatic gradients (both 4.5h and 39h) and the expression of each gene across the 200 ROIs of the left brain based on Schaefer-400 atlas. The significance of the PLS was tested again using a permutation analysis (n = 10,000). It finally derived a vector of 15631 correlation coefficients representing the degree of certain gene spatially aligns with the pattern of glymphatic gradient.

#### Gene ontology (GO) analysis

GO analysis was performed using Metascape platform^28^ for gene lists positively or negatively associated with the PLS-derived transcriptomic profiles. We reported the false discovery rate (FDR)-adjusted significance values generated by Metascape using the Benjamini-Hochberg procedure across the tested enrichment terms within each analysis.

#### Gene set enrichment analysis (GSEA)

We further performed GSEA on the whole gene list (i.e., the ordered set of 15631 genes from PLS analysis) using abagen^18, 27^ to investigate the transcriptomic profile of 30 brain cell-types (astrocytes, endothelial cells, pericytes, microglia, oligodendrocytes, oligodendrocytes progenitor cells, 13 subtypes of excitatory neurons, and 11 subtypes of inhibitory neurons) related with glymphatic gradient. For each brain cell gene set, an enrichment score representing the level of enrichment was obtained. To correct for the size of different gene sets, the enrichment score was compared with those estimated from permutation tests (n = 10,000) and derived a normalized enrichment score (NES) for each gene set. FDR values were taken directly from the GSEA workflow in abagen, without an additional multiple-testing adjustment.

### Cortical Aβ map

The Aβ map was originally collected by Ranasinghe et al.^19^ and was downloaded from the NeuroVault (https://neurovault.org/). Briefly, PET was acquired with PiB to visualize Aβ deposits in the brain of 32 cognitively impaired patients with Alzheimer’s disease (mild cognitive impairment or dementia stage). Individual SUVR maps were computed in native space using data acquired between 50 and 70 min after tracer injection, and using the cerebellum gray matter as a reference region (defined using FreeSurfer 5.3). The 32 SUVR maps were finally averaged across patients. The average SUVR map for PiB was parcellated based on the Schaefer-400 atlas.

### Definition of fALFF- or ReHo-glymphatic clearance match-mismatch pattern

First, mean fALFF, ReHo and 39h PC maps were calculated across individuals. Then, the mean fALFF, ReHo and 39h PC across ROIs were transformed to z-scores, respectively. FALFF-glymphatic clearance match pattern was defined as ROIs with z-fALFF > 0 and z-39h PC < 0, or z-fALFF < 0 and z-39h PC > 0. FALFF-glymphatic clearance mismatch pattern was defined as ROIs with z-fALFF > 0 and z-39h PC > 0 (mismatch1), or z-fALFF < 0 and z-39h PC < 0 (mismatch2). Same definition applied to ReHo-glymphatic clearance match, mismatch1, and mismatch2 patterns. For each ROI, fALFF-glymphatic clearance mismatch index was defined as the negative absolute difference between z-fALFF and z-39h PC (i.e., –|z-fALFF – z-39hPC|). Higher score indicated less difference between z-fALFF and z-39h PC, indicating higher neural activity with slower clearance or lower neural activity with faster clearance, i.e., greater mismatch between fALFF and 39h PC. ReHo-glymphatic clearance mismatch index was defined likewise, i.e., –|z-ReHo – z-39h PC|, with higher scores indicating greater mismatch between ReHo and 39h PC.

### Statistical analyses

Statistical analyses were performed in SPSS packages (IBM, Chicago, version 22.0 for Windows) and MATLAB R2020a (MathWorks, Inc.). We checked the normality of all the characteristics by using the Kolmogorov-Smirnov test and histogram inspection. Statistical significance was accepted at the 0.05 level (two-tailed).

#### Comparison of PC

One-way repeated measure one-way analysis of variance (ANOVA) was used to compare regional PC across timepoints with post-hoc analyses. Independent two-sample t tests were used to compare regional PC between patients with neurodegenerative disease and peripheral neuropathy. Paired two-sample t test was used to compare the mean PC of patients with neurodegenerative disease and peripheral neuropathy across ROIs. Pearson correlations were used to investigate the association of PC between patients within different disease subgroups, and to assess accordance of gradient scores with PC in 4.5h or 39h across ROIs with spin test to adjust for spatial auto-correlation^29^.

#### Association among fALFF, 39h PC, fALFF-39h PC mismatch index, and Aβ burden

The primary correlations involving fALFF, 39h PC, fALFF-39h PC mismatch index, and Aβ burden were calculated across cortical parcels using group-level maps. Parcel counts denote the spatial units entering a correlation. Pearson correlation between fALFF and 39h PC was tested with spin tests. Independent ANOVA was used to compare the Aβ burden among match, mismatch1, and mismatch2 regions with FDR-controlled multiple comparisons (i.e., two-stage linear step-up procedure of Benjamini, Krieger and Yekutieli). Furthermore, Spearman correlations were used to investigate the associations among z scores of fALFF and 39h PC, mismatch index, and Aβ burden in match, mismatch1, and mismatch2 regions. In association between mismatch index and Aβ burden, partial Spearman correlations were used when adjusting for z-scores of fALFF and 39h PC, and FDR-corrected. The Benjamini-Hochberg procedure was applied separately to each tests across the match, mismatch1, and mismatch2 regions. Uncertainty in the correlation coefficients was estimated using 10,000 paired participant-level bootstrap resamples.

#### Sensitivity analysis

To confirm the robustness of the fALFF-39h PC match and mismatch regions, a K-means clustering algorithm was programmed in Python. First, the fALFF and 39h PC were standardized using z-score normalization to mitigate scale-related biases. Then, the K-means clustering used K = 4 to determine if the clusters were concordant with those defined in fALFF-39h PC mismatch index. Since K-means clustering does not provide specific meaning, the clusters were manually assigned as distinct patterns in reference to clusters in fALFF-39h PC mismatch index. To evaluate the robustness of the group-level fALFF-39 h PC spatial correlation to the omission of individual participants, we performed a leave-one-out analysis in the rs-fMRI subgroup (n = 15). Each participant was omitted in turn, and the group-mean fALFF and 39 h PC maps were recalculated from the remaining 14 participants. Pearson’s correlation was then recomputed across the 400 cortical parcels.To replicate the results regarding neural activity based on fALFF, ReHo and ReHo-39h PC mismatch index were analyzed. The analysis procedure and statistics replicated those involving fALFF.

## Results

### Participant Characteristics

Following selection flowchart (Supplementary Fig. 1), 96 participants were included to characterize the cortical glymphatic function (57 ± 14 [SD] years; 49 women). Disease diagnosis is listed in Supplementary Table 1. Among them, 15 participants without intracranial diseases and with rs-fMRI were further included to investigate the association between spontaneous neural activity and cortical glymphatic function (53 ± 12 [SD] years; 8 women), of whom 11 were diagnosed as peripheral neuropathy and 4 as motor neuron disease. Demographic and clinical information are listed in Table 1.

**Table 1.**
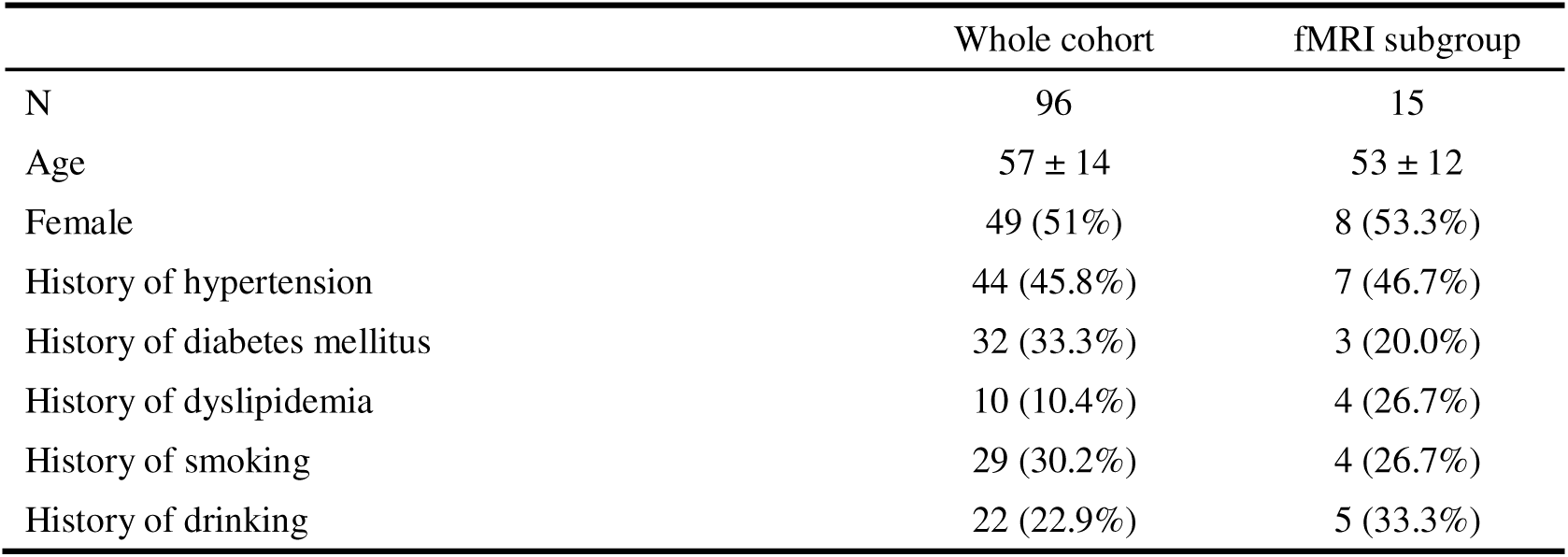
Population characteristics.

|  | Whole cohort | fMRI subgroup |
| --- | --- | --- |
| N | 96 | 15 |
| Age | 57 ± 14 | 53 ± 12 |
| Female | 49 (51%) | 8 (53.3%) |
| History of hypertension | 44 (45.8%) | 7 (46.7%) |
| History of diabetes mellitus | 32 (33.3%) | 3 (20.0%) |
| History of dyslipidemia | 10 (10.4%) | 4 (26.7%) |
| History of smoking | 29 (30.2%) | 4 (26.7%) |
| History of drinking | 22 (22.9%) | 5 (33.3%) |

### Glymphatic dynamics across cortex

Regional glymphatic dynamics and their comparisons between different timepoints are illustrated in Fig. 2. For 4.5h, medial prefrontal cortex and insula were among regions with fast glymphatic influx, while temporal lobe was the slowest (Fig 2A). For 39h, periventricular regions like cingulate cortex, temporal lobe, and occipital lobe were with fast glymphatic clearance, while surface regions like dorsal prefrontal and parietal lobe were slower (Fig 2C). At individual level, PC significantly differed across timepoints (P < 0.001, Fig 2F). Overall, both at regional and individual levels, PC increased from 4.5h to 15h, then decreased to 39h. Thus, higher 4.5h PC could indicate better glymphatic influx, while higher 39h PC indicates worse glymphatic clearance.

**Fig. 2.**
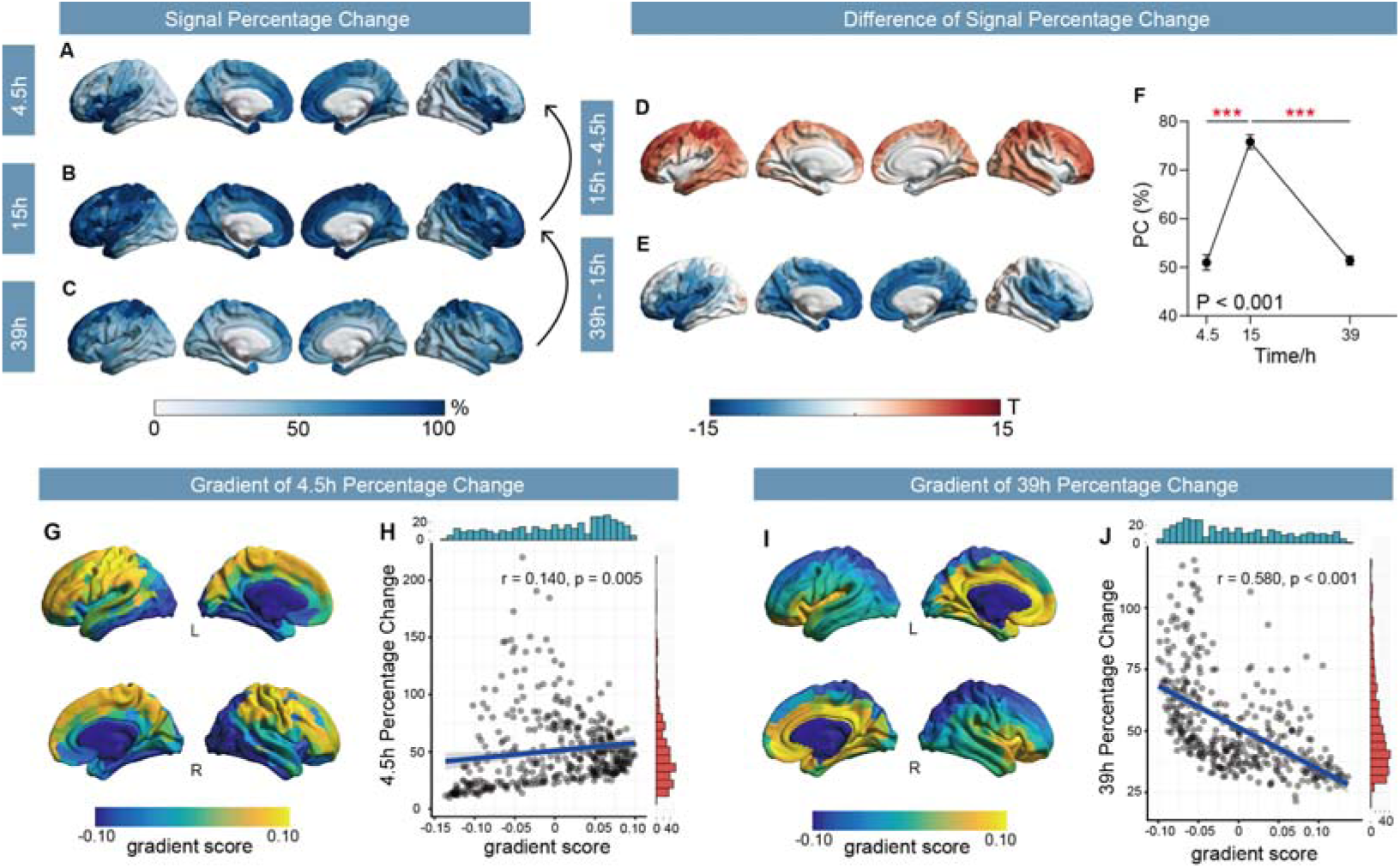
Regional glymphatic dynamics across the cortex. (A-C) Glymphatic signal percentage change at 4.5h (A), 15h (B), and 39h (C) across cortex based on Schaefer-400 atlas. (D-E) T statistics from paired t-tests comparing the signal percentage change between 15h and 4.5h (D), as well as 39h and 15h (E) across cortex. (F) Line graph of 4.5, 15, and 39h percentage change (PC) in 400 ROIs across cortex. Dot represents mean, and bar represents standard error of the mean. (G, I) The first gradient map across cortex based on Schaefer-400 atlas for 4.5h (G) and 39h percentage change (I). The scatter plots show Pearson correlations between first gradient score and percentage change at 4.5h (H) and 39h (J). The translucent area around the regression lines represents the 95% confidential interval for the regression estimate, with fit line (blue) for significant association. The blue histogram represents the distribution of variables in x-axis, whereas the red histogram represents the distribution of variables in y-axis. L, left; R, right. ***, p < 0.001.

While previous studies have widely posited that glymphatic transport is reduced in neurodegenerative diseases, it remains unclear whether this represents a uniform slowing with preserved spatial relationships between brain regions, or a more fundamental reorganization of the overall clearance pattern. Surprisingly, the general distribution patterns of CSF tracer were similar and highly correlated (Fig. 3K, M) between neurodegenerative disease (n = 17) and peripheral neuropathy (n = 49, served as control) subgroups across timepoints (Fig. 3A-F). On a group-level, pair-wise comparison of mean PC across 400 ROIs showed that neurodegenerative disease patients may have a generally delayed glymphatic influx and clearance than those with peripheral neuropathy (Fig. 3J, L). In ROI-wise analysis, neurodegenerative disease patients had lower 4.5h PC in frontal cortex than peripheral neuropathy. Besides, patients with neurodegenerative diseases tended to exhibit higher PC in the cingulate cortex compared to those with peripheral neuropathy from 4.5h to 39h. By 39 hours, this trend had expanded to temporal lobe and frontal cortex; however, none of the observed differences survived FDR correction (Fig. 3G–I).

**Fig. 3.**
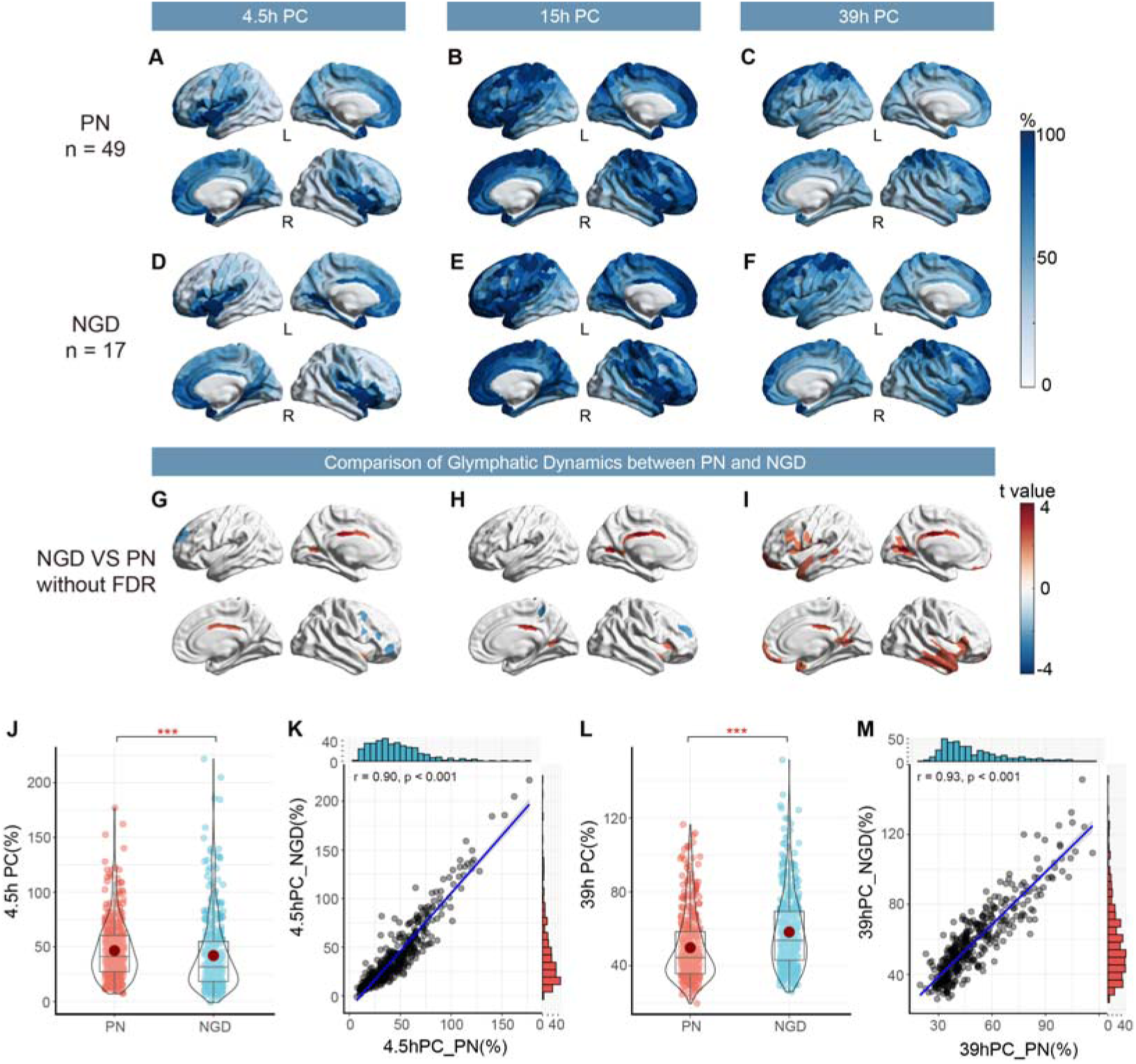
Comparison of glymphatic dynamics between patients with peripheral neuropathy (PN) and neurodegenerative diseases (NGD). (A-C) Glymphatic signal percentage change (PC) at 4.5h, 15h, and 39h in PN (n = 49). (D-F) Glymphatic signal PC at 4.5h, 15h, and 39h in NGD (n = 17). (G-I) T statistics from independent two sample t tests of PC at 4.5h, 15h, and 39h between NGD and PN without false discovery rate (FDR) correction. (J) Violin plot of paired two sample t test of 4.5h PC between PN and NGD. (K) Scatter plot of Pearson correlation of 4.5h PC between PN and NGD. (L) Violin plot of paired two sample t test of 39h PC between PN and NGD. (M) Scatter plot of Pearson correlation of 39h PC between PN and NGD. In K and M, the translucent area around the regression lines represents the 95% confidential interval for the regression estimate, with fit line (blue) for significant association. The blue histogram represents the distribution of variables in x-axis, whereas the red histogram represents the distribution of variables in y-axis. L, left; R, right. ***, p < 0.001.

### Cortical glymphatic gradient

To capture the intrinsic property of cortical glymphatic dynamics, gradients were calculated for 4.5h PC and 39h PC across cortex, respectively (Fig. 2G, I). In specific, gradient was positively correlated with PC value at 4.5h (Fig. 2H), while negatively correlated with PC value at 39h (Fig. 2J). Generally, the influx gradient follows an anterior-posterior hierarchy. Meanwhile, the clearance gradient follows a hierarchy from periventricular to surface area.

### Transcriptomic correlates of glymphatic influx and clearance

To explore the potential molecular foundation underlying these macroscopic imaging patterns, we performed a spatial transcriptomic analysis leveraging AHBA. Using GO analysis with 4.5h PC gradient, we found several pathways enriched in regions with faster influx, broadly mapping to gene regulation and synapse function, such as “chromatin remodeling” (Count of genes = 113, -log10(P) = 17.4), “cell junction organization” (Count of genes = 99, -log10(P) = 17.3), “DNA repair” (Count of genes = 120, -log10(P) = 14.0), “regulation of synapse structure or activity” (Count of genes = 76, -log10(P) = 12.8) (Fig. 4A, B). Meanwhile, regions with slower influx enriched pathways related with metabolism, such as “aerobic respiration and respiratory electron transport” (Count of genes = 497, -log10(P) = 70.7), “mitochondrion organization” (Count of genes = 127, -log10(P) = 45.8), “mitochondrial protein degradation” (Count of genes = 47, -log10(P) = 25.6). (Fig. 4C, D).

**Fig. 4.**
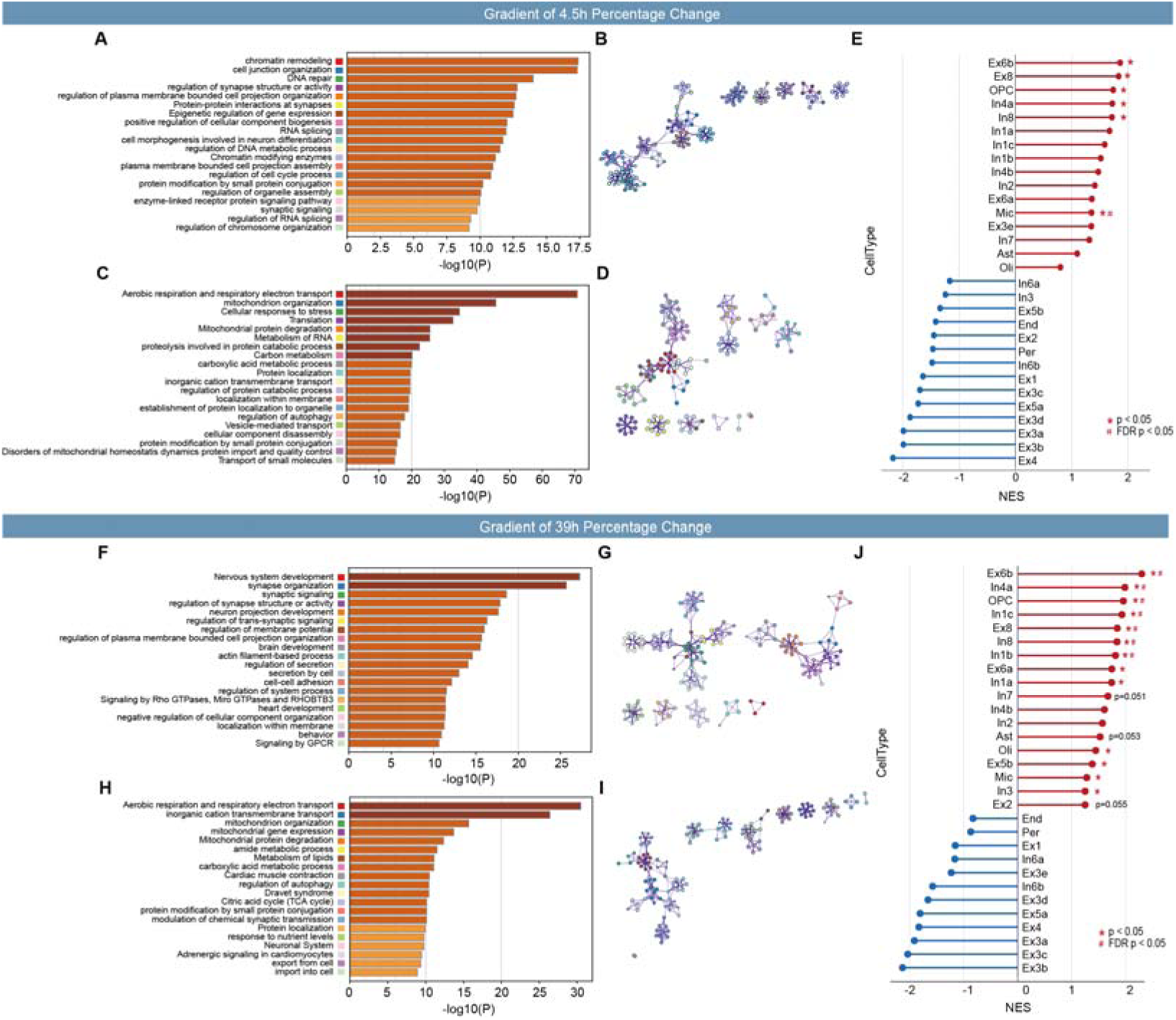
Association between first gradient of glymphatic flow and gene expression profiles. (A-E) shows the result for the first gradient of 4.5h percentage change. (A) The top 20 GO biological processes linked with gene sets positively correlated with the first gradient of 4.5h percentage change, and (B) shows the gene enrichment network. (C) The top 20 GO biological processes linked with gene sets negatively correlated with the first gradient of 4.5h percentage change, and (D) shows the gene enrichment network. (E) Cell type enrichment analysis of the first gradient map. (F-J) shows the result for the first gradient of 39h percentage change. (F) The top 20 GO biological processes linked with gene sets positively correlated with the first gradient of 39h percentage change, and (G) shows the gene enrichment network. (H) The top 20 GO biological processes linked with gene sets negatively correlated with the first gradient of 39h percentage change, and (I) shows the gene enrichment network. (J) Cell type enrichment analysis of the first gradient map. Colors indicate directions of association (red = positive, blue = negative). *, p < 0.05; #, false discovery rate (FDR) p < 0.05.

Similarly, in 39h PC gradient, regions with faster clearance enriched pathways related with synapse function, such as “synapse organization” (Count of genes = 107, -log10(P) = 25.7), “synaptic signaling” (Count of genes = 135, -log10(P) = 18.6), “regulation of synapse structure or activity” (Count of genes = 85, -log10(P) = 17.9) (Fig. 4F, G). Regions with slower clearance enriched pathways related with metabolism, such as “aerobic respiration and respiratory electron transport” (Count of genes = 385, -log10(P) = 30.5), “inorganic cation transmembrane transport” (Count of genes = 209, -log10(P) = 26.4), “amide metabolic process” (Count of genes = 92, -log10(P) = 11.5). (Fig. 4H, I).

Next, we assessed whether the genes associated with glymphatic dynamics, were strongly expressed in specific cell types. GSEA analysis found that, oligodendrocyte precursor cell (NES = 2.00, p = 0.004, FDR p = 0.030), astrocyte (NES = 1.77, p = 0.012, FDR p = 0.06), oligodendrocyte (NES = 1.62, p = 0.002, FDR p = 0.030), microglia (NES = 1.61, p < 0.001, FDR p < 0.001), and some types of inhibitory neurons (for example, In1c, NES = 1.65, p = 0.004, FDR p = 0.030) were enriched in regions with faster influx (Fig. 4E). As for clearance, both excitatory (for example, Ex6b, NES = 2.30, p < 0.001, FDR p < 0.001) and inhibitory (for example, In1c, NES = 1.93, p < 0.001, FDR p < 0.001) neurons, oligodendrocyte precursor cell (NES = 1.95, p < 0.001, FDR p < 0.001), oligodendrocyte (NES = 1.44, p = 0.044, FDR p = 0.01), microglia (NES = 1.28, p = 0.006, FDR p = 0.022) were enriched in regions with faster clearance (Fig. 4J). Astrocyte was enriched with a tendency (NES = 1.52, p = 0.053). Overall, above results showed a close relationship of synaptic structure and function with intact glymphatic function, as well as metabolism with impaired glymphatic function, triggering us to investigate the association between spontaneous neural activity and glymphatic function.

### Association between spontaneous neural activity and glymphatic clearance across cortex

The cortical 39h PC of participants with fMRI (n = 15) was highly coherent with that of the whole cohort (Fig. 5A-B). Across 400 ROIs in cortex, z-score of fALFF was negatively correlated with 39h PC ( r = -0.20, 95% CI [-0.350, -0.007]; Fig. 5C-D), suggesting that impaired glymphatic clearance often coupled with lower neural activity. The group-level spatial correlation between fALFF and 39 h PC remained negative across all leave-one-out analyses (r range, -0.239 to -0.161), indicating that its direction was not driven by any single participant.

**Fig. 5.**
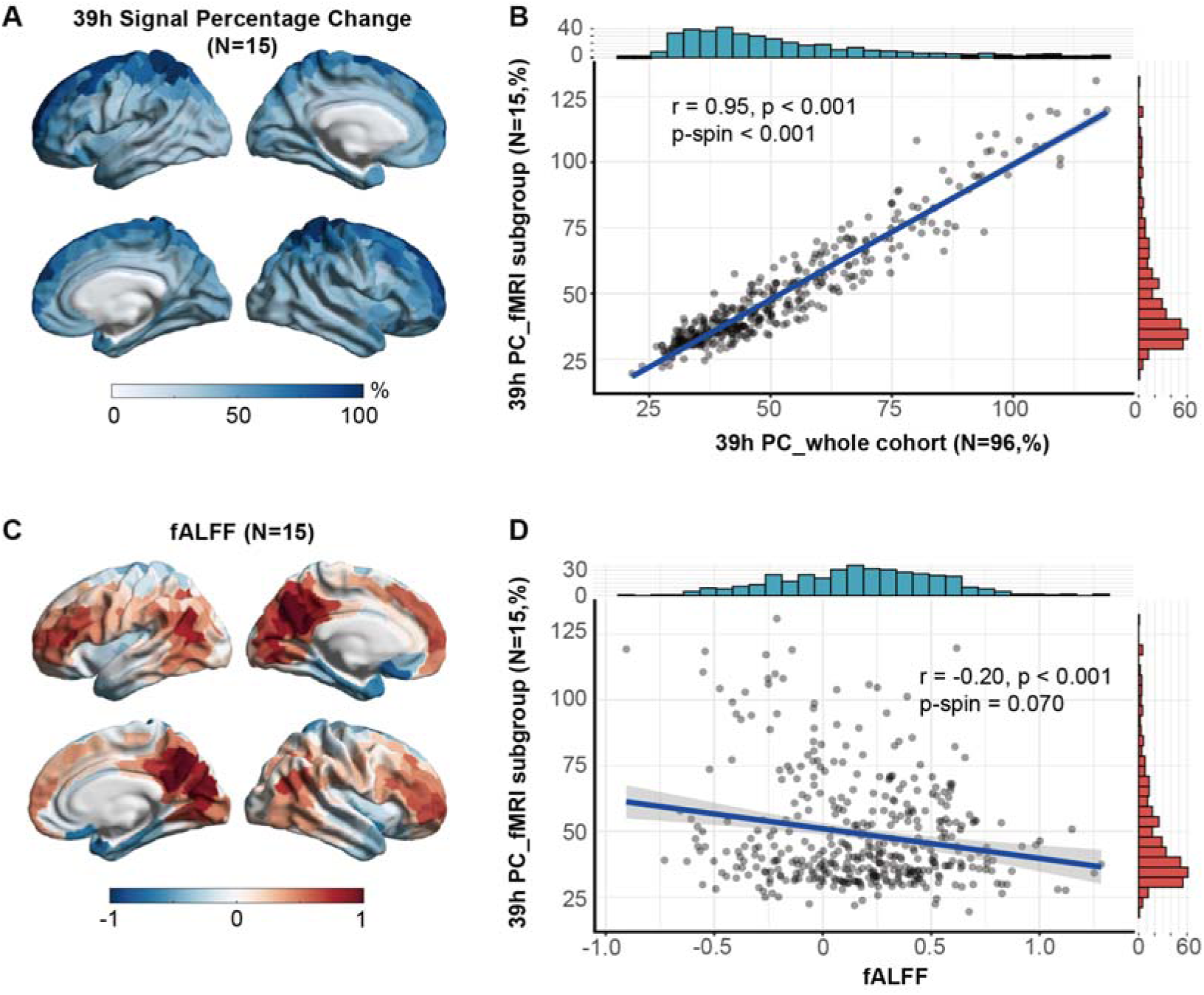
Association between fractional ALFF (fALFF) and 39h percentage change (PC) across cortex. (A) The 39h PC across cortex based on Schaefer-400 atlas in patients with fMRI (n = 15). (B) The scatter plot of Pearson correlation between 39h PC in whole cohort (n = 96) and subgroup with fMRI (n = 15). (C) The fALFF across cortex based on Schaefer-400 atlas in patients with fMRI (n = 15). (D) The scatter plot of Pearson correlation between fALFF and 39h PC and across cortex based on Schaefer-400 atlas in patients with fMRI (n = 15). In B and D, the translucent area around the regression lines represents the 95% confidential interval for the regression estimate, with fit line (blue) for significant association. The blue histogram represents the distribution of variables in x-axis, whereas the red histogram represents the distribution of variables in y-axis.

### Cortical fALFF-glymphatic clearance mismatch pattern overlaps with amyloid-prone cortical regions

Match regions mainly located in parietal and occipital lobes, while mismatch1 regions (high activity-slow clearance) in dorsal prefrontal cortex, and mismatch2 regions (low activity-fast clearance) in anterior cingulate cortex and insula (Fig. 6A). Data-driven clustering revealed a similar pattern, further confirmed its robustness (Supplementary Fig. 2). Utilizing open-source ^11^C-PiB PET SUVR dataset^19^ (Fig. 6B), we found fALFF-39h PC mismatch regions were more overlapped with higher Aβ burden than match regions (Fig. 6C, one-way ANOVA F = 4.391, p = 0.013; Post-hoc: mismatch1 vs match, p = 0.031; mismatch2 vs match, p = 0.010).

**Fig. 6.**
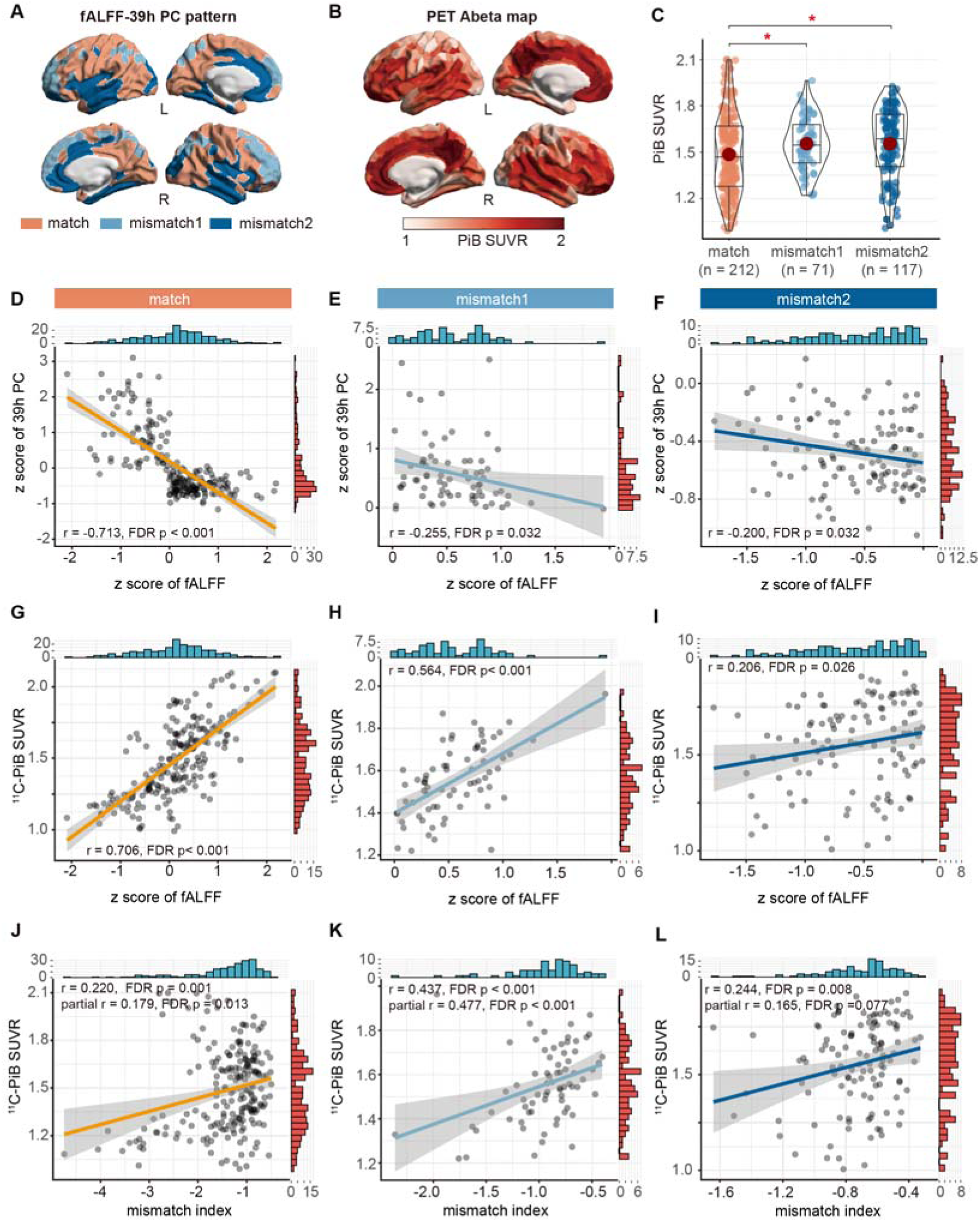
Association between fALFF-39h percentage change (PC) mismatch and amyloid-β (Aβ) burden. (A) fALFF-39h PC match (orange), mismatch1 (high activity with low clearance, light blue) and mismatch2 (low activity with high clearance, dark blue) regions. (B) Open-source ^11^C-PiB SUVR for Aβ PET imaging based on Schaefer-400 atlas. (C) Independent one-way ANOVA test of PiB SUVR among regions in match (orange), mismatch1 (light blue) and mismatch2 (dark blue) pattern. p values are calculated using FDR-controlled multiple comparison: two-stage linear step-up procedure of Benjamini, Krieger and Yekutieli. (D-F) The scatter plot of Spearman correlation between z score of fALFF and 39h PC in match (D), mismatch1 (E) and mismatch2 (F) regions. (G-I) The scatter plot of Spearman correlation between z score of fALFF and PiB SUVR in match (G), mismatch1 (H) and mismatch2 (I) regions. (J-L) The scatter plot of Spearman correlation between fALFF-39h PC mismatch index and PiB SUVR in match (J), mismatch1 (K) and mismatch2 (L) regions. In D-L, the translucent area around the regression lines represents the 95% confidential interval for the regression estimate, with fit line for significant association; r indicates the Spearman correlation coefficient, and partial r indicates the Spearman partial correlation coefficient when adjusting for z scores of fALFF and 39h PC;the Benjamini-Hochberg procedure was applied separately to each tests across the match, mismatch1, and mismatch2 regions. FDR, false discovery rate. *, p < 0.05. L, left; R, right.

In line with definition, with z-score of fALFF increasing, match regions showed decreased z-score of 39h PC, indicating better clearance; while mismatch regions showed weaker associations with z-score of 39h PC (match regions, r= -0.713, 95% CI [-0.735, -0.635]); mismatch1 regions, r = -0.255, 95% CI [-0.421, -0.046]); mismatch2 regions, r = -0.200, 95% CI [-0.297, -0.001]; Fig. 6D-F).

In match regions, z-score of fALFF was positively correlated with Aβ burden (Fig. 6E), but weaker in mismatch regions ( match regions, r= 0.706, 95% CI [0.629, 0.743]); mismatch1 regions, r = 0.564, 95% CI [0.328, 0.658]); mismatch2 regions, r= 0.206, 95% CI [-0.013, 0.316]; Fig. 6G-I).

Across cortex, identified mismatch regions overlap with amyloid-prone cortical regions after adjustment for z-scores of fALFF and 39h PC, which was stronger in mismatch1 regions than in match regions or mismatch2 regions. Both match and mismatch1 regions survived adjustment for z-scores of fALFF and 39h PC, leaving mismatch2 regions with a tendency (match regions, r=0.179, 95% CI [0.056, 0.266]; mismatch1 regions, r =0.477, 95% CI [0.166, 0.560]; mismatch2 regions, r =0.165, 95% CI [−0.051, 0.328]; Fig. 6J-L).

### Sensitivity Analysis

Clustering analysis showed great similarities between the ReHo-39h PC pattern and fALFF-39h PC pattern (Supplementary Fig. 3A-B), with their mismatch index strongly correlated (Supplementary Fig. 3C).

Z-score of ReHo was negatively correlated with z-score of 39h PC. Notably, in mismatch2 regions, this association was insignificant (match regions, r= -0.639, 95% CI [-0.703, -0.554]; mismatch1 regions, r=-0.236, 95% CI [-0.400, -0.002]; mismatch2 regions, r=-0.092, 95% CI [-0.170, 0.088]; Supplementary Fig. 3D-F). Consistent with fALFF, z-score of ReHo was positively correlated with Aβ burden in match regions (match regions, r= 0.609, 95% CI [0.488, 0.676]; Supplementary Fig. 3E), though weaker in mismatch regions (mismatch1 regions, r =0.491, 95% CI [0.220, 0.543]; mismatch2 regions, r= 0.194, 95% CI [0.016, 0.271]; Supplementary Fig. 3G-I). Across cortex, regional ReHo-39h PC mismatch index was positively correlated with Aβ burden after adjustment for z-scores of ReHo and 39h PC, which was stronger in mismatch1 regions than in match regions or mismatch2 regions (match regions, r=0.279, 95% CI [0.150, 0.369]; mismatch1 regions, r = 0.274, 95% CI [0.012, 0.435]; mismatch2 regions, r =0.260, 95% CI [0.094, 0.342]; Supplementary Fig. 3J-L).

## Discussion

Capitalizing on Glymphatic MRI, this study deciphered the tripartite interplay among glymphatic dynamics, neural activity, and amyloidosis. At the molecular level, neuroimaging-transcriptional analyses on the glymphatic pattern demonstrated that genes associated with synapse organization and activity were significantly enriched in brain regions with better glymphatic influx and clearance. At the cellular level, the linked genes were predominantly expressed in specific cell types, including excitatory or inhibitory neurons, and several glial cells. At the regional level, based on a subgroup of participants with rs-fMRI, we demonstrated that glymphatic clearance was positively correlated with spontaneous neural activity. Mismatch index was defined to evaluate decoupling between spontaneous neural activity and glymphatic clearance function, which turned out that identified mismatch regions largely overlap with amyloid-prone cortical regions.

Our systematic mapping of cortical glymphatic dynamics revealed a conserved spatiotemporal architecture, demonstrating notable interindividual consistency. Overall, glymphatic fluid flows in at 4.5h and clears at 39h, aligning with previous studies^30^. To decode the underlying cortical hierarchy^31, 32^, we derived distinct gradients for cortical glymphatic influx and clearance function. Generally, the influx gradient follows an anterior-posterior hierarchy, orchestrating with the distribution of gamma frequency band^33^, a neurophysiological signature previously linked to glymphatic potentiation^12^. Meanwhile, the clearance gradient follows a hierarchy from periventricular to surface area, leaving questions on how extracellular diffusion or advection contribute to this process. Notably, though coherent in pattern distribution, patients with neurodegenerative diseases exhibited regionally impaired glymphatic clearance capability compared with peripheral neuropathy controls. This deficit was particularly prominent in regions predisposed to Aβ aggregation^1, 34^ like cingulate and frontal cortex, reinforcing the hypothesized role of glymphatic failure as a neurodegeneration accelerant^35^. Together, our results present an intrinsically stable glymphatic dynamic pattern. Elucidating its structural basis will advance understanding of fluid dynamics and clearance processes in the brain.

Evidence for coupling between neuronal activity and glymphatic function remains scarce in human. Utilizing AHBA dataset, we demonstrated that the transcriptomic profiles of 10 excitatory or inhibitory neuronal subtypes were positively correlated with glymphatic clearance function, with 4 subtypes enriched in regions with better glymphatic influx. Zooming in, we also found that spatial variation in glymphatic influx and clearance mirrored variation in neural activity-related molecular pathways, particularly genes governing synaptic architecture and plasticity. Strikingly, glymphatic clearance was associated with spontaneous neural activity including activity amplitude (fALFF) and synchronization (ReHo). Our results, together with other studies^11, 13, 36^, identifying an important yet overlooked player functionally coupled with glymphatic flow beside vascular pulsations: the neurons. Neurons can drive parenchymal convection flow. Experimental models demonstrate that coordinated neuronal firing generates interstitial ionic waves capable of enhancing cerebrospinal fluid perfusion and clearance^11^, while neuromodulatory interventions, including transcranial optogenetics in rodents^11^ or visual stimulation in humans^36^, markedly amplify clearance efficacy. Intriguingly, neurons can also regulate vasomotion through distant neurovascular coupling^13^, or by regulating vasoactive intestinal peptide interneurons through non-invasive 40Hz stimulation^12^. Recently, Zhang et al demonstrated that glutamatergic axons dilate innervating arterioles via synaptic-like transmission at neural-arteriolar smooth muscle cell junctions^37^. This synaptic-like structure not only provide novel insights in communication between neurons and hemodynamic fluctuation, but also offer a potential structural basis for loop between neurons, vasomotion and perivascular glymphatic dynamics. Future research is required to delineate the intricate interactions between neurons and glymphatic system.

Importantly, spatial heterogeneity was observed in the relationship between neuronal activity and glymphatic function. Using novel fALFF- or ReHo-39hPC mismatch index and data-driven k-means clustering, a relatively robust activity-clearance pattern was observed. Notably, match regions localized predominantly to parietal and occipital lobes, mismatch1 regions to dorsal prefrontal cortex, and mismatch2 regions to anterior cingulate cortex and insula. Previously, based on ADNI data with evaluation of CSF movement in the fourth ventricle, Han et al creatively depicted the coupling between regional BOLD signal and CSF^38^. Regions with higher coupling in their findings align substantially with our match pattern located in motor/sensory, auditory and visual networks, which mostly belong to the lower-order regions. Meanwhile, both mismatch1 and mismatch2 localized to regions capable of higher-order cognition^39^, yet how mismatch influence their functioning remains elusive. The detailed biological mechanisms driving or influencing mismatch still needs further investigations.

Activity-clearance mismatch represents a novel spatial correlate associated with regional proteopathy in the brain. Intriguingly, we discovered that both mismatch1 and mismatch2 regions exhibited higher Aβ burden than match regions. Moreover, the mismatch index was generally positively associated with Aβ burden after adjustment for neural activity and glymphatic clearance, especially in mismatch1 regions. While both neural activity and glymphatic clearance dysfunction have been implicated in Aβ decomposition^3, 35, 40^, their cooperative role in Aβ accumulation remains understudied. Notably, Aβ spreads preferentially through neural pathways^41, 42^, not fully compatible with the direction of glymphatic flow. This paradox may be partially reconciled considering the mismatch mechanism, suggesting the hypothesis that the imbalance between neural activity and glymphatic function may affect the final spreading pattern of toxic proteins. Previously, the coupling between resting-state global activity and CSF movement has been found associated with neocortical tau deposition^43^ and various AD-related pathology. While informative, these studies only evaluate CSF movement in the fourth ventricle but not glymphatic flow in parenchyma, with regional variability largely ignored. Our quantitative studies based on Glymphatic MRI shed light on this question by detailing in vivo assessment of the mismatch between regional glymphatic clearance and neuronal activity. At the preclinical stage of Alzheimer disease, Aβ mainly accumulates in default mode network and, to a lesser degree, the frontoparietal network^34^, overlapping partially with our mismatch1 pattern. It is also worth noting that brain regions showing age-invariant glucose metabolism are more susceptible to AD-related hypometabolism^4^ and overlap with our mismatch2 pattern, potentially involving activity-clearance mismatch. Whether augmenting glymphatic clearance function could mitigate mismatch and alleviate subsequent proteopathy in neurodegenerative diseases remains a question for future research.

Astrocytes are indispensable components in glymphatic system as CSF flushes along the perivascular spaces formed by their end-feet, with deletion of astrocyte AQP4 resulting in waste deposition^5^. In our cell type analysis, astrocyte enriched in regions with faster glymphatic influx and tended to enrich in regions with faster clearance, partially supporting its role in glymphatic maintenance. Except for the well-known astrocytes, better glymphatic clearance function was found correlated with genes expressed in other glia cells including oligodendrocyte, oligodendrocyte progenitor cell, and microglia. Typically, oligodendrocytes myelinate axons and provide metabolic support to neurons^44^; microglia provide immune surveillance, synaptic pruning, and debris removal^45^. Anatomically, using state-of-the-art technics, Allen et al confirmed that oligodendrocytes were enriched in the corpus callosum or near the CSF^46^. In ageing, glial aging was particularly accelerated in white matter compared with cortical regions^47^, while white matter serves as a preferential, low-resistance glymphatic pathway under normal conditions^6, 48^. Meanwhile, aging-induced activation of microglia showed a strong dependence on their proximity to oligodendrocytes^46^. Together, above evidence suggest oligodendrocyte and microglia as vulnerable foci when glymphatic system impairs. Consistently, in human, glymphatic dysfunction is closely related with demyelination diseases including white matter hyperintensity^49^ and multiple sclerosis^50^. It is plausible that glymphatic clearance impairment promotes accumulation of inflammatory and neurotoxic elements, activates microglia, and disrupts oligodendrocyte lineage determination, ultimately exacerbates demyelination. These results encourage future investigations on the mechanism between glymphatic function and other neuroglia.

Several issues warrant consideration. First, our participants were with different diseases, though a stable glymphatic dynamic pattern was established. Further studies exploring disease-related effects are needed. Meanwhile, neuroinflammation may contribute to prolonged retention of gadodiamide, which could delay the cortical signal enhancement, inducing potential bias. Future comparisons using macrocyclic agents or validated non-gadolinium tracers may help distinguish altered tracer transport from tissue-related retention. Second, only three time points after intrathecal injection were obtained, leaving relatively long period of blank. Third, since glymphatic dynamic is under circadian control^35, 40^ and influenced by posture^51^, scan time (though set relatively fixed), individual sleep-wake cycle, sleeping posture or day-time activity may constitute potential confounders. Furthermore, regional variations in cortical thickness, adjacent CSF volumes, tissue integrity and vascular density may alter the local PC signal independent of true parenchymal tracer penetration, causing partial volume effect. Residual confounding by age, sex, diagnosis, vascular risk factors, renal function, and other clinical characteristics remains possible. Fourth, it is crucial to distinguish between measured MRI signal dynamics and the actual microscopic glymphatic transport mechanisms. Due to the spatial resolution limits of standard MRI, the delayed tracer retention (proxy for impaired clearance) observed in our study may stem from diminished convective driving forces, altered interstitial diffusion barriers, or a combination of both. Fifth, the cross-sectional design precludes causal inferences regarding glymphatic-neural activity mismatch and Aβ deposition. Sixthly, transcriptomic analyses relied solely on the AHBA data in left hemisphere, precluding lateralization assessment. As the gene expression profiles derive from healthy participants, our conclusions reflect intrinsic variability rather than disease-associated changes in transcriptomics. Seventhly, because the amyloid burden map was derived from an external open-source dataset, the observed association represents a spatial correspondence across group-averaged maps rather than an individual-level relationship. Given the cross-sectional design and the limited fMRI sample size (n=15), the directionality and exact temporal sequence between activity-clearance mismatch and proteopathy cannot be established. This relationship is likely complex and modulated by age and the stage of neurodegeneration. Future longitudinal studies with concurrent PET, fMRI, and glymphatic MRI are warranted to elucidate these temporal dynamics. Lastly, this study mainly focused on general relationship between neural activity and glymphatic clearance across regions, neglecting the possibility that hub regions in network may have different properties.

In conclusion, rather than demonstrating a generic interplay, our study specifies that macroscopic cortical regions experiencing activity-clearance mismatch exhibit profound spatial overlap with amyloid-prone territories. Our imaging proxy map for glymphatic clearance could be analyzed integrative with a series of cerebral molecular, physiological, and cognitive-related features^33, 52^, potentially advancing our understanding towards the intricate role of glymphatic system in multimodal and multiscale neuroscience. Further investigations are necessary to determine the cellular and molecular constituents in interaction between neurons and glymphatic system, and how they are affected by diseases.

## Acknowledgements

This study was supported by the National Natural Science Foundation of China (82171276, U23A20426) and Research Project on Prevention and Treatment Techniques for Stroke of Development Center for Medical Science & Technology, National Health Commission of the People’s Republic of China (WKZX2023CZ0220).

## Conflicts of Interest

The authors declare no competing interests.

## Author Contributions

Y.L., X.Z., and M.L. conceptualized and designed the study; Y.L., X.Z., Y.Z., X.Z., Z.Z., K.W., and J.S. collected the data; Y.L. and X.Z. analyzed the data and prepared the figures; Y.L. and X.Z. wrote the original manuscript draft; Y.Z. and M.L. reviewed and edited the manuscript; M.L. supervised the study. All authors have read and approved the final version of this manuscript.

## Data sharing statement

The datasets used and/or analyzed during the current study are available from the corresponding author on reasonable request.

## Supplementary Material

**Supplementary Table 1.**

| Diagnosis | N (%) |
| --- | --- |
| Peripheral neuropathy | 49 (51.0%) |
| Motor neuron disease | 8 (8.3%) |
| Encephalitis | 16 (16.7%) |
| Suspected CSF leakage | 3 (3.1%) |
| Hepatic encephalopathy | 1 (1.0%) |
| Multiple sclerosis | 2 (2.1%) |
| Neurodegenerative diseases | 17 (17.7%) |
| Normal pressure hydrocephalus | 5 (5.2%) |
| Parkinson's disease | 4 (4.2%) |
| Possible cerebral amyloid angiopathy | 3 (3.1%) |
| Multiple system atrophy | 3 (3.1%) |
| Alzheimer's disease | 1 (1.0%) |
| Lewy body dementia | 1 (1.0%) |

**Supplementary Figure 1.**
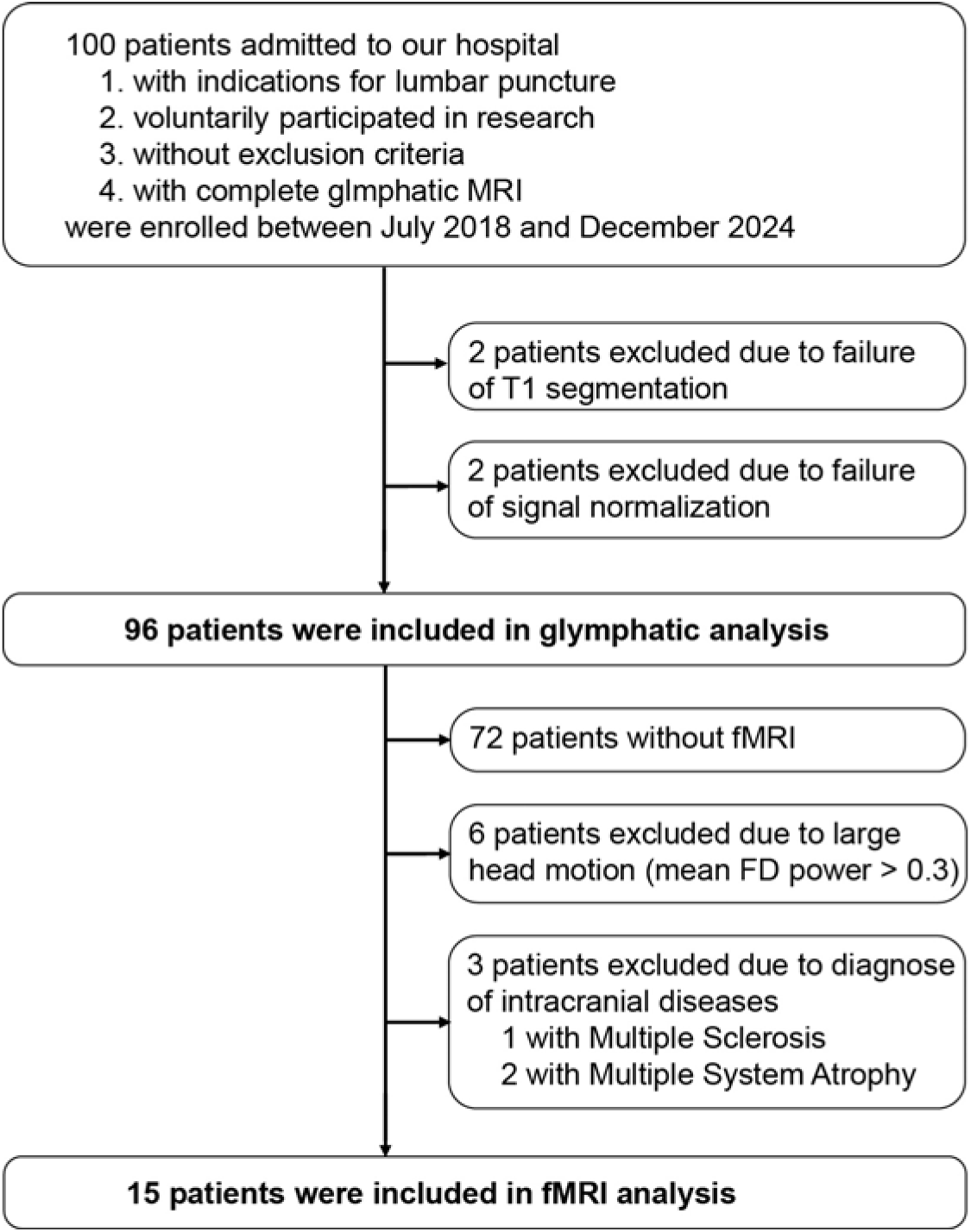
Flowchart of patient selection.

**Supplementary Figure 2.**
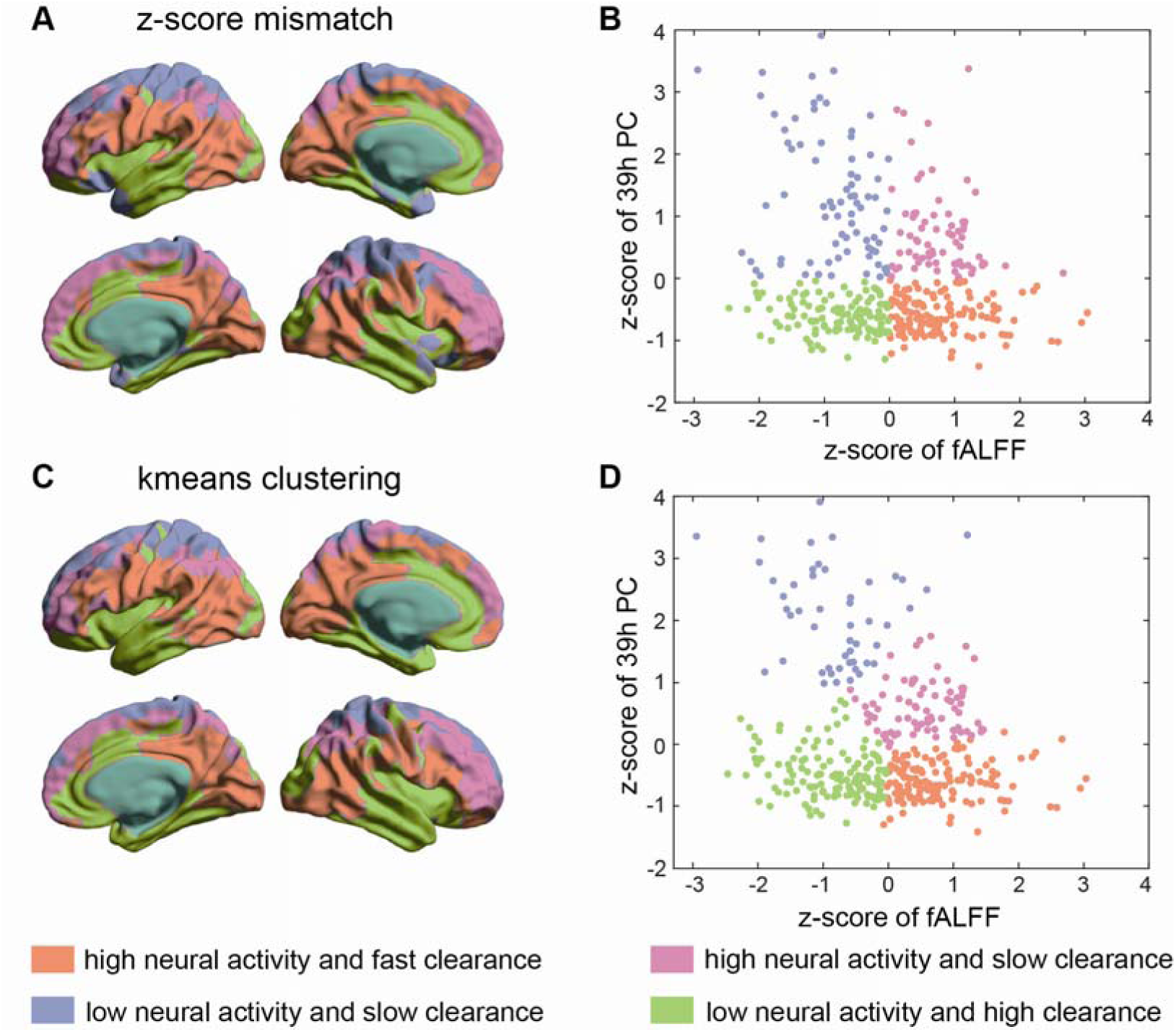
fALFF-39h percentage change (PC) pattern based on different clustering methodology. (A-B) The spatial location (A) and scatter plot (B) of clusters based on fALFF-39h PC mismatch index. (C-D) The spatial location (C) and scatter plot (D) of data-driven k-means clustering based on z-scores of fALFF and 39h PC. Orange, high neural activity and fast clearance; pink, high neural activity but slow clearance; purple, low neural activity and slow clearance; green, low neural activity but fast clearance.

**Supplementary Figure 3.**
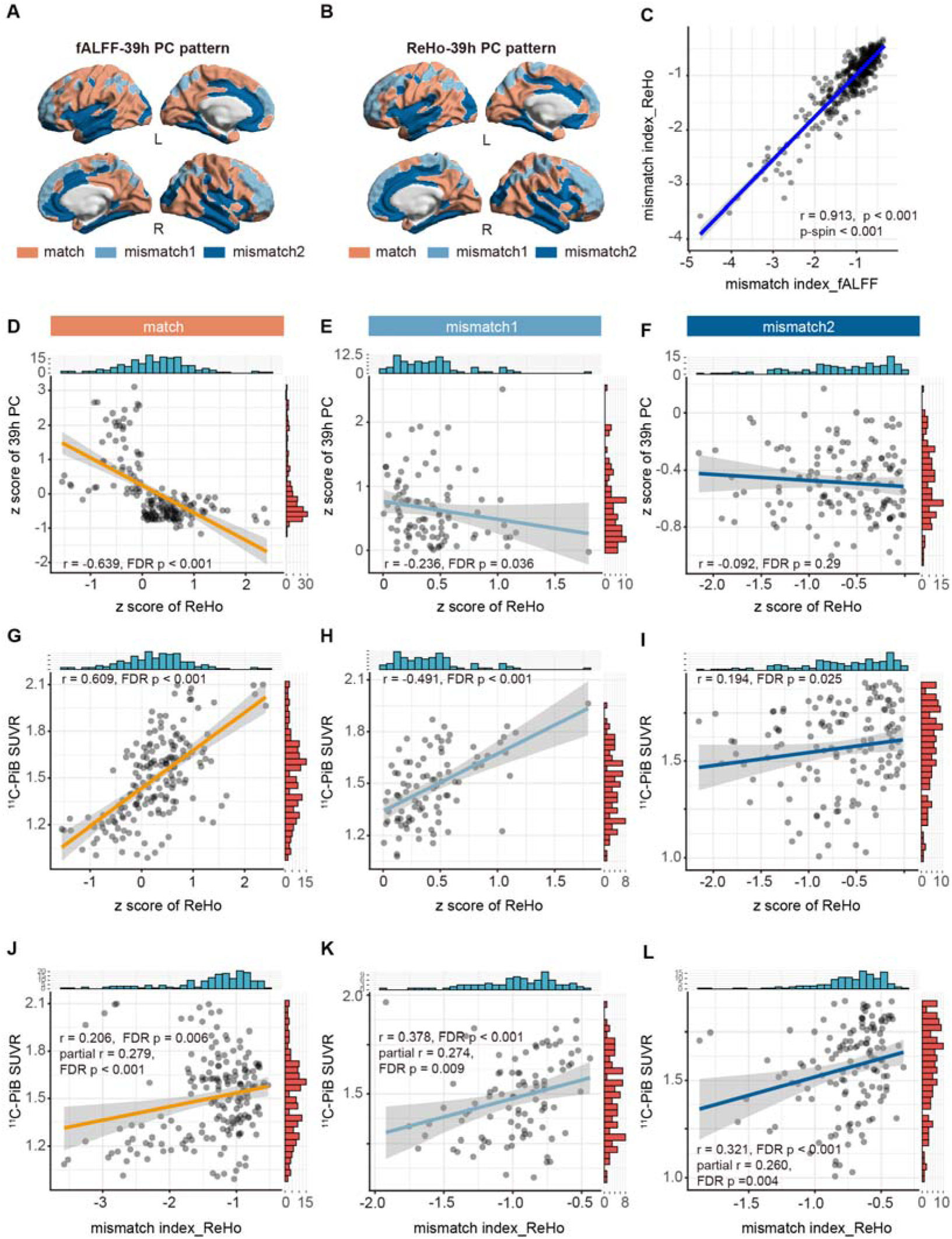
Association between ReHo-39h percentage change (PC) mismatch and amyloid-β (Aβ) burden. (A) fALFF-39h PC match (orange), mismatch1 (high activity with slow clearance, light blue) and mismatch2 (low activity with fast clearance, dark blue) regions. (B) ReHo-39h PC match (orange), mismatch1 (high activity with slow clearance, light blue) and mismatch2 (low activity with fast clearance, dark blue) regions. (C) The scatter plot of Pearson correlation between fALFF-39h PC mismatch index and ReHo-39h PC mismatch index. (D-F) The scatter plots of Spearman correlation between z score of ReHo and 39h PC in match (D), mismatch1 (E) and mismatch2 (F) regions. (G-I) The scatter plots of Spearman correlation between z score of ReHo and PiB SUVR in match (G), mismatch1 (H) and mismatch2 (I) regions. (J-L) The scatter plots of Spearman correlation between ReHo-39h PC mismatch index and PiB SUVR in match (J), mismatch1 (K) and mismatch2 (L) regions. In D-L, the translucent area around the regression lines represents the 95% confidential interval for the regression estimate, with fit line for significant association; r indicates the Spearman correlation coefficient, and partial r indicates the Spearman partial correlation coefficient when adjusting for z scores of ReHo and 39h PC; the Benjamini-Hochberg procedure was applied separately to each tests across the match, mismatch1, and mismatch2 regions. FDR, false discovery rate. L, left; R, right.

## Notes

### Competing Interest Statement

The authors have declared no competing interest.

### Summary of Updates

1. Clarifying the imaging measure. We distinguish cortical tracer mapping in 96 participants from the activity-clearance analyses in the 15 participants with rs-fMRI. The Methods state: Conversely, the PC at the delayed time point (39 h) represents prolonged tracer retention, serving as a proxy for impaired glymphatic clearance. Our interpretation therefore reflects the indirect nature of this measure. We have replaced assertive terms like direct glymphatic flow with precise descriptors such as imaging proxies for tracer enhancement/retention. Furthermore, we expand the Limitations section to explicitly acknowledge the confounding effects of Partial Volume Averaging (PVE), systemic tracer redistribution, renal clearance kinetics, and so on. 2. Assessing robustness in the fMRI subgroup. We performed complementary leave-one-out analyses. Notably, these analyses show that the negative group-average fALFF-39 h PC estimate is not dependent on single participant. We have thoroughly updated Limitation paragraph in Discussion. 3. Improving statistical reporting. We have expanded reporting of effect sizes, confidence intervals, and multiple-comparison procedures. We distinguish participant-sampling uncertainty from spatial inference and clarify that associations with the external amyloid map represent regional spatial correspondence rather than relationships across individuals. 4. Aligning conclusions with the evidence. We have revised the conclusions to emphasize spatial associations and acknowledge the limitations of causal interpretation.

